# Enhancing Generative Decoders with Stochastic Training on Biomedical Data with Missingness

**DOI:** 10.64898/2025.12.01.690409

**Authors:** Xi Shen, Andreas Bjerregaard, Yan Li, Anders Krogh

## Abstract

Biomedical datasets are often heterogeneous and affected by noise or missing values. Deep Generative Decoders (DGD) provide a promising framework for latent representation learning, but their standard training procedure relies on sample-level stochastic gradient descent (SGD), which performs poorly with incomplete data. To address this, we introduce two stochastic training strategies — Nested SGD and Feature Dropout — that incorporate feature-level randomness into optimization. Evaluations on biomedical tabular datasets demonstrate that Nested SGD improves robustness under missingness the best, while Feature Dropout not only improves but also accelerates convergence with lower computational cost. These results suggest that feature-level stochasticity is a practical way to strengthen biomedical AI pipelines.

## 1 Introduction

Biomedical research increasingly relies on data-driven methods to interpret complex and heterogeneous datasets [LeCun et al., 2015]. However, challenges such as incomplete patient records, measurement noise, and variability across cohorts often limit the effectiveness of conventional representation learning approaches. The Deep Generative Decoder (DGD) has emerged as an alternative to autoencoders [Schuster and Krogh, 2023], directly optimizing per-sample latent vectors for reconstruction (Fig. 1A). While variational autoencoders are inherently challenged by input with missing variables (requiring non-trivial fixes [Collier et al., 2020]), the DGD relies only on observed features. This makes it a natural fit to handle data with missingness. Recent developments also highlight its generalizability in manifold learning and Riemannian representations [Bjerregaard et al., 2025]. While effective, DGD typically uses SGD [Bottou, 2012, Hardt et al., 2016], which performs poorly and struggles to generalize under incomplete data [Little and Rubin, 2019, Nazábal et al., 2020].

**Figure 1:**
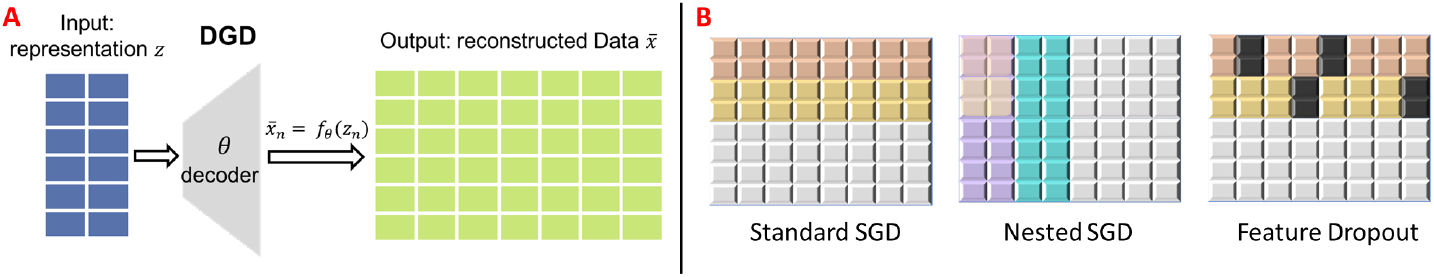
A) Model schematic: DGD optimizes a representation of each sample with standard back-propagation. B) Training strategies: Standard SGD, Nested SGD, Feature Dropout. Rows represent samples and columns represent features.

To overcome these limitations, we introduce **feature-level stochastic training** for DGDs: (1) Nested SGD and (2) Feature Dropout. We keep the DGD encoder-free and use stochastic *feature subsampling* only in the loss, optimizing per-sample latents on the observed entries. Unlike masked-token prediction [Devlin et al., 2018] and denoising autoencoders [Vincent et al., 2008] — which corrupt **x** and train an encoder–decoder to reconstruct masked or corrupted entries— our method never imputes or cleans missing features. On three biomedical tabular datasets, these strategies improve robustness under missingness; Nested SGD is most accurate, Feature Dropout is most efficient.

## 2 Methods and Experiments

In standard SGD, stochasticity arises from computing gradients on minibatches of samples., which introduces noise during training. In the DGD setting, however, each sample’s latent representation receives only a single gradient update per epoch, making it less affected by this noise. To increase stochasticity at the representation level, we therefore propose to also sample features stochastically. This is feasible because the model contains no encoder and the reconstruction loss naturally ignores missing features. The differences between standard SGD and the proposed methods are illustrated in Fig. 1B. Pseudocode is provided in Appendix A.

### Nested SGD

Each training step samples both data instances and feature subsets, introducing dual-level stochasticity. We assume that this regularizes learning and improves robustness at the expense of higher runtime.

### Feature Dropout

Random subsets of features are excluded from the loss computation, encouraging resilience to missingness and reducing reliance on any single feature [Li et al., 2019]. We assume this approach is computationally efficient but less stable.

### Datasets and Evaluation

We evaluate on three tabular datasets covering diverse biomedical applications: Thyroid Disease [Yadav, 2020], Diabetes Health Indicators [UCI Machine Learning Repository, 2015], and Parkinson Telemonitoring [UCI Machine Learning Repository, 2009]. Fully connected DGDs are applied to these datasets. Artificial missingness is introduced under the Missing Completely at Random (MCAR) assumption [Rubin, 1976]; details of the masking procedure and the masked reconstruction loss are given in Appendix B.

**Table 1:** Overview of datasets used in evaluation.

| Dataset | Samples × Features | Natural Missing | Feature Type | Task |
| --- | --- | --- | --- | --- |
| Thyroid Disease | 9 172 × 31 | Yes | Multivariate | Classification |
| Parkinson Telemonitoring | 5 875 × 19 | No | Continuous | Regression |
| Diabetes | 21 207 × 21 | No | Multivariate | Classification |

Three different experiments were conducted to evaluate model behavior under different patterns of missingness in training and testing; an overview is shown in Table 2. The experiments measure downstream utility (random forest regression or classification of a relevant target) as missingness varies, see Appendix C. Optimization hyperparameters for each method and dataset, such as learning rate, weight decay and betas, are provided in Appendix D.

**Table 2:**
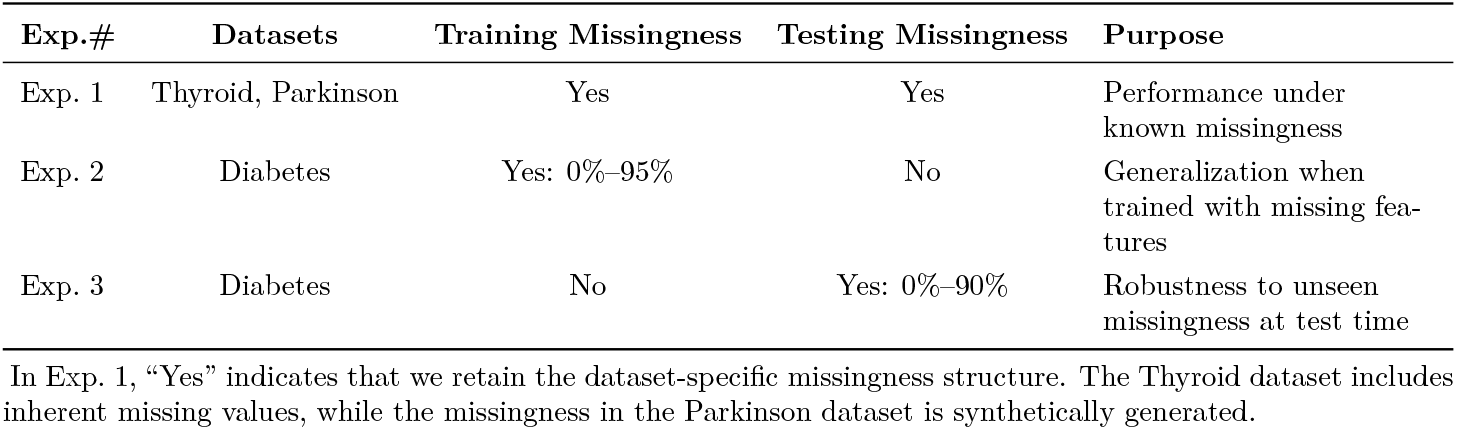
Overview of the three experiments.

## 3 Results

In Experiment 1, Fig. 2A, the F1-score of the Thyroid dataset improved from 0.759 (Standard SGD) to 0.799 (Nested SGD) and 0.802 (Feature Dropout). For the Parkinson regression task, the test MSE decreased from 47.8 (Standard SGD) to 45.4 (Nested SGD) and 44.8 (Feature Dropout), as shown in Fig. 2B. These results demonstrate that introducing feature-level stochasticity leads to more reliable representations for downstream biomedical prediction tasks. The performance gains are also more pronounced on the Thyroid dataset, which happens to exhibit real missingness and strong class imbalance.

**Figure 2:**
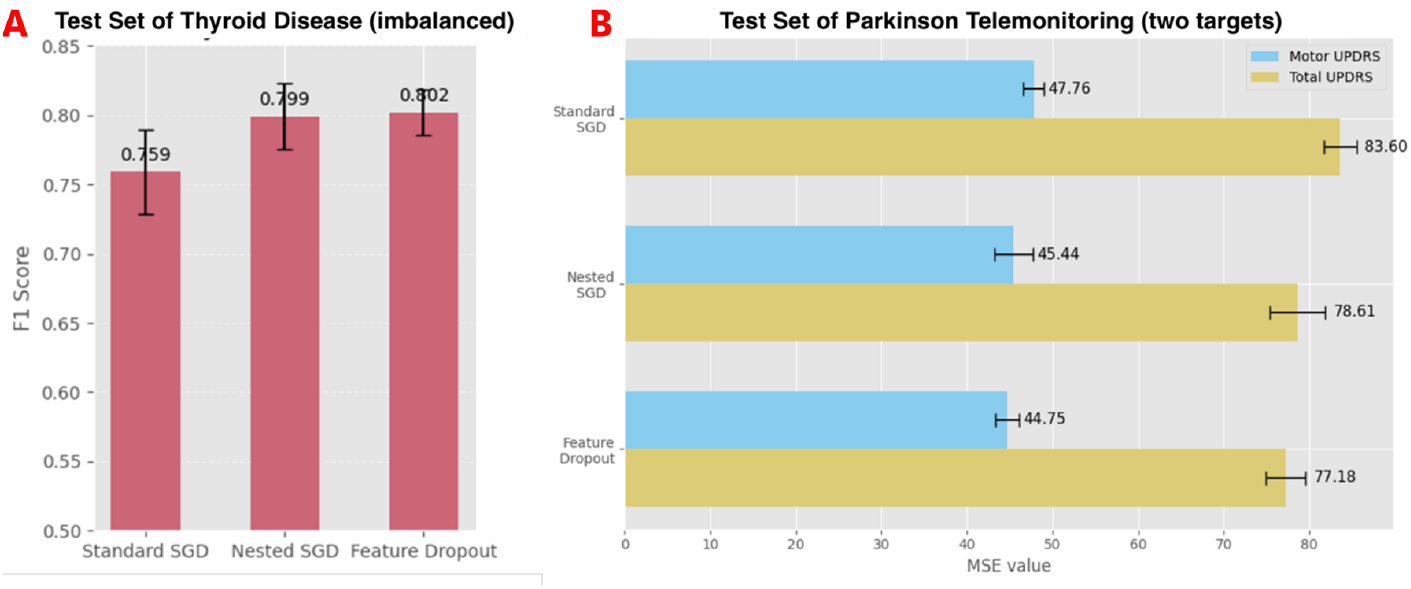
Results of Experiment 1. (A) F1-score of the Thyroid Disease dataset. (B) MSE of the Parkinson Telemonitoring dataset.

In Experiment 2, the test accuracy curves in Fig. 3A show that both Nested SGD and Feature Dropout achieve consistently stronger generalization than Standard SGD, with the advantage becoming more pronounced under higher levels of training missingness. Among the three methods, Nested SGD attains the highest and most stable accuracy, with Feature Dropout yielding closely comparable performance. The selected training loss curves (Fig. 3B and C) further illustrate that both methods converge in fewer epochs than Standard SGD under representative missingness levels.

**Figure 3:**
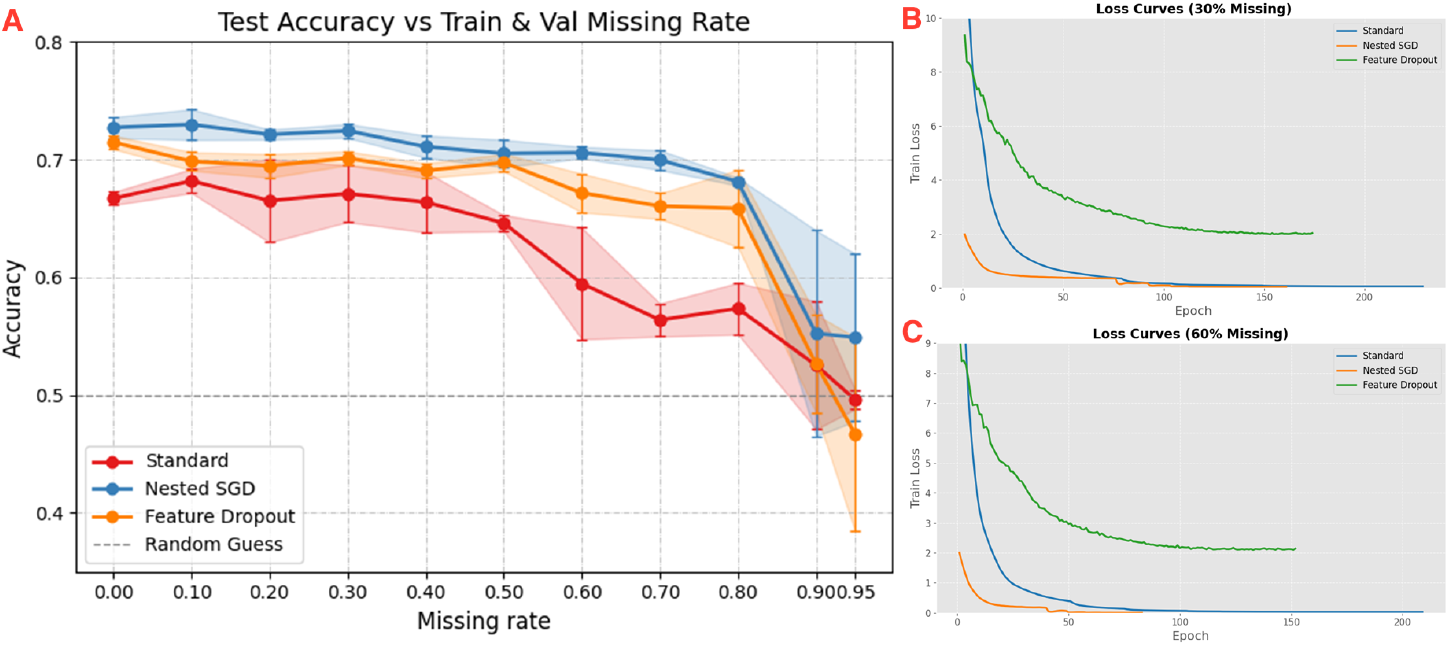
Results of Experiment 2 (saturating missingness during *training*). (A) Test accuracy on the diabetes dataset as *training missingness* increases from 0% to 95%; (B, C) Training loss curves under 30% and 60% missingness.

Experiment 3 (Fig. 4) follows the same setup as Experiment 2 but introduces missingness only during testing. As shown in Fig. 4A, Nested SGD again achieves the highest and most stable accuracy across different levels of test-time missingness, followed by Feature Dropout. The loss curves in Fig. 4B and C show that Feature Dropout converges more quickly than the other methods, but with noticeably noisier optimization trajectories. Taken together, these results indicate that the three approaches offer different trade-offs that may be relevant in practice, depending on whether robustness to test-time missingness or computational efficiency is prioritized.

**Figure 4:**
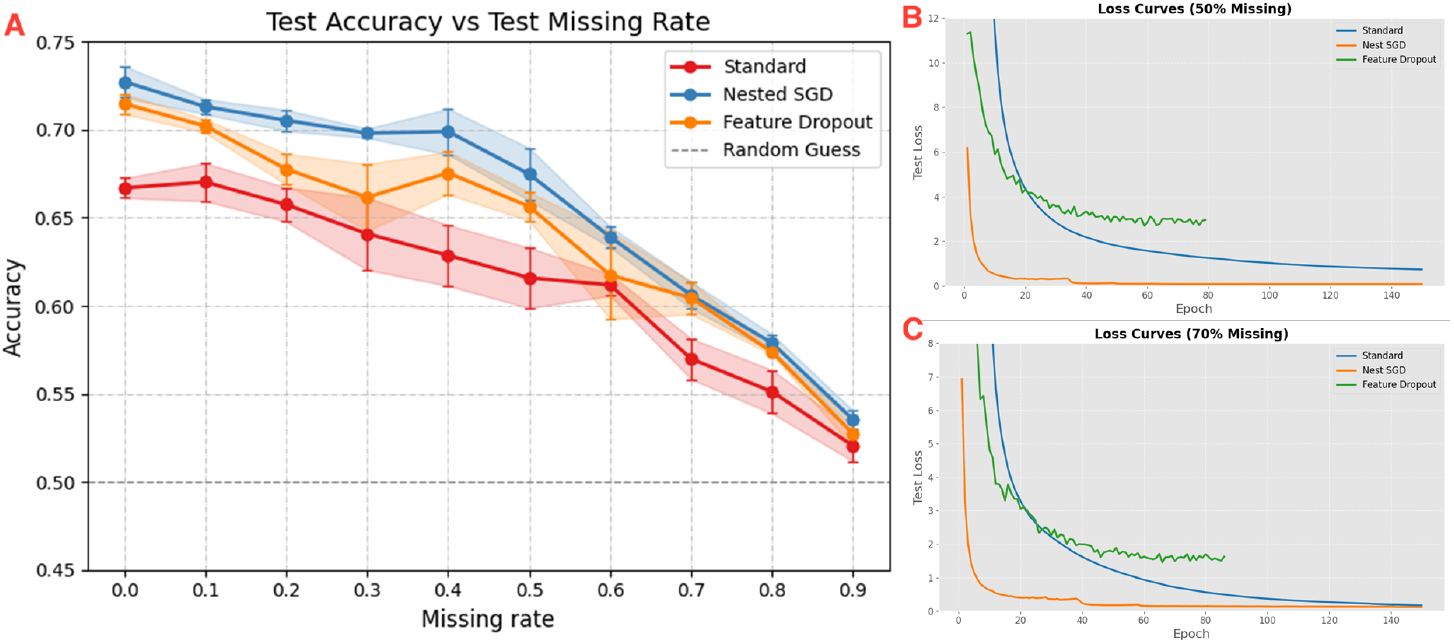
Results of Experiment 3 (saturating missingness during *testing*). (A) Test accuracy on the diabetes dataset as *test missingness* increases from 0% to 90%; (B, C) Test loss curves under 50% and 70% missingness.

Finally, Table 3 summarizes the average runtime for every 10 epochs across datasets and methods. While Nested SGD offers improved robustness, it introduces higher computational overhead because of its nested optimization loop. In contrast, Feature Dropout achieves the best efficiency among all methods. Note that the absolute runtime values depend on hardware and GPU configuration; the relative comparison across methods, however, remains consistent.

**Table 3:**
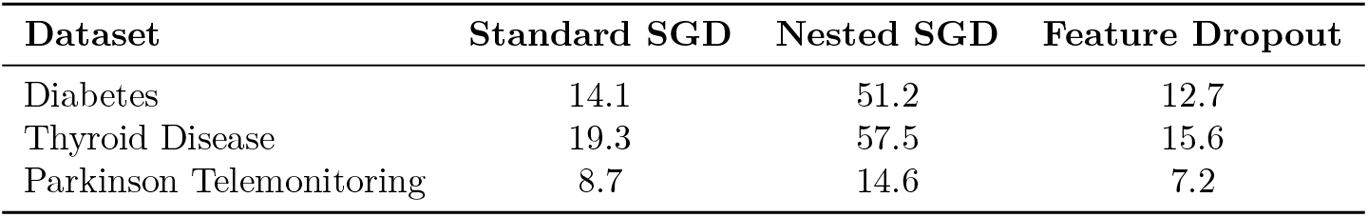
Average runtime for every 10 epochs (seconds) across methods.

## 4 Conclusions

This study demonstrates that both Nested SGD and Feature Dropout can enhance generalization and robustness under incomplete biomedical data, with Nested SGD showing overall superior performance across tasks. These results suggest that introducing feature-level stochasticity not only improves the predictive accuracy of DGDs but also facilitates convergence under missingness. However, each approach has its limitations: Nested SGD increases computational cost due to its two-level optimization structure, while Feature Dropout may exhibit instability depending on the dropout rate and data sparsity.

Future work can extend to more intricate missingness mechanisms, explore data with higher underlying dimensionality, and apply more intricate decoder architectures. These steps would further enhance the practical relevance of this work on biomedical data, where systematic biases in data collection are common. We also envision collaborations with biomedical scientists to integrate these stochastic strategies into pipelines for disease prediction and longitudinal cohort studies.

## Author Contributions

X.S. designed the study, collected the datasets, developed the algorithms, implemented the framework, performed the analyses, and wrote the manuscript. A.B., Y.L. and A.K. supervised the study and contributed to conceptual development, methodology and manuscript revision.

## Acknowledgements

This work was supported by grants NNF20OC0062606 and NNF23OC0083510 from the Novo Nordisk Foundation. AK was further supported by grants NNF20OC0063268 and NNF20OC0059939.

## A Algorithms of Stochastic Training Strategies

### Algorithm 1 Nested SGD Training

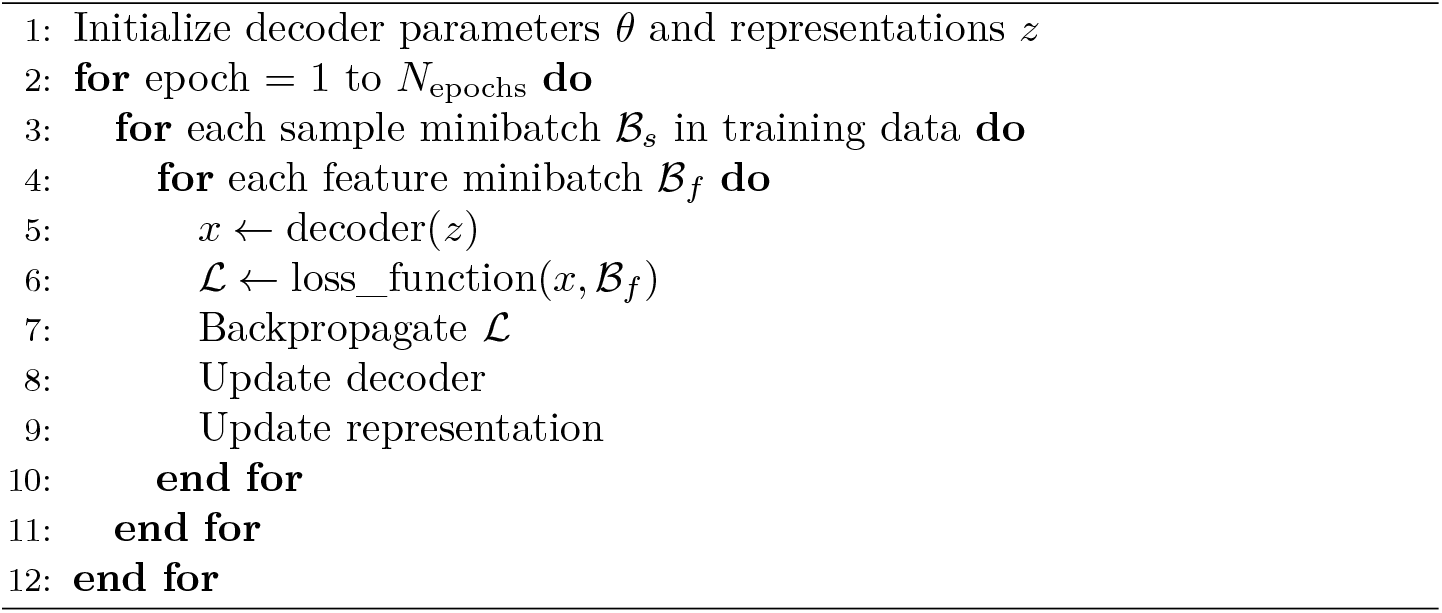

### Algorithm 2

Feature Dropout Training

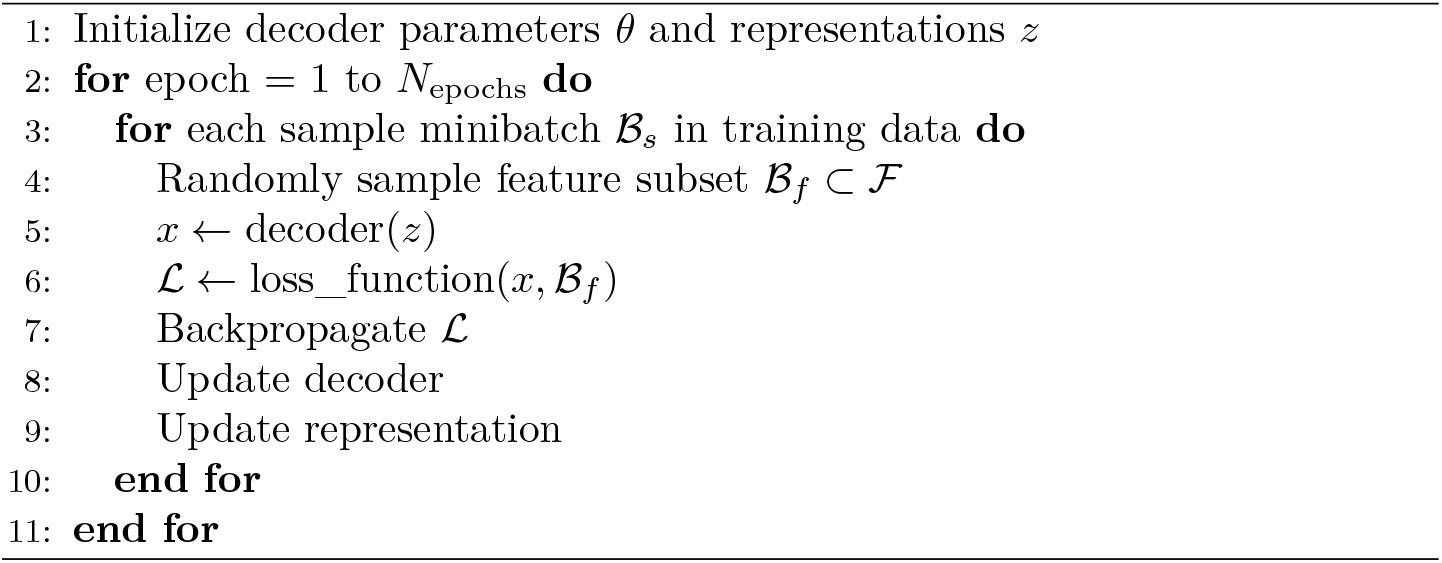

## B Details of Missingness Handling and Masked Reconstruction Loss

To handle incomplete data and to artificially induce MCAR missingness, we follow the general masking strategy outlined in Nazábal et al. [2020], but with a simplified implementation. Missing or to-be-masked values are replaced with zeros in the input **x**_*n*_, but the decoder is not explicitly given the binary mask. Instead, the mask is used only to restrict the reconstruction loss to observed (non-missing) features.

### B.1 Mask Definition

Let **x**_*n*_ = (*x*_*n*1_, …, *x*_*nD*_) be the feature vector for sample *n*. The binary mask **m**_*n*_ = (*m*_*n*1_, …, *m*_*nD*_) is defined as:

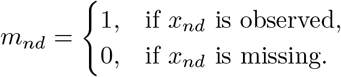

The reconstruction loss for a single sample is:

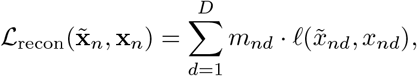

where *l*(·) denotes a pointwise loss (e.g., squared error). The total loss is obtained by summing over all samples.

### B.2 Heterogeneous Feature Types

Continuous features are standardized, categorical features are one-hot encoded, and binary features are kept unchanged. We split inputs and masks into three groups:

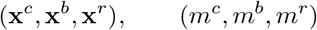

where *c, b*, and *r* denote categorical, binary, and real-valued features.

### B.2 Masked Likelihood Loss

Let 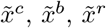 be reconstructed outputs. The overall loss is a weighted sum of masked negative log-likelihoods:

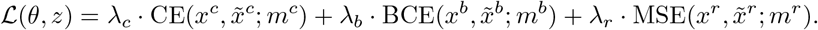

Each component is defined as:

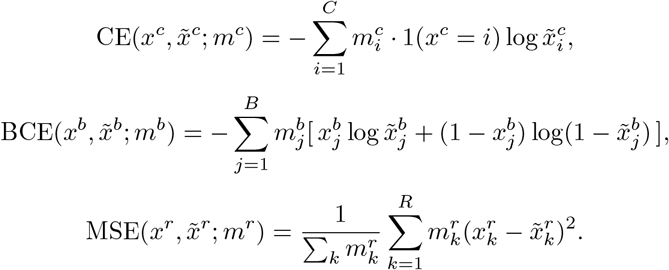

Only the continuous (MSE) term is normalized by the number of observed values, as MSE scales with variance and dimensionality; CE/BCE are already normalized by construction.

## C Experiment Pipeline

**Figure S1:**
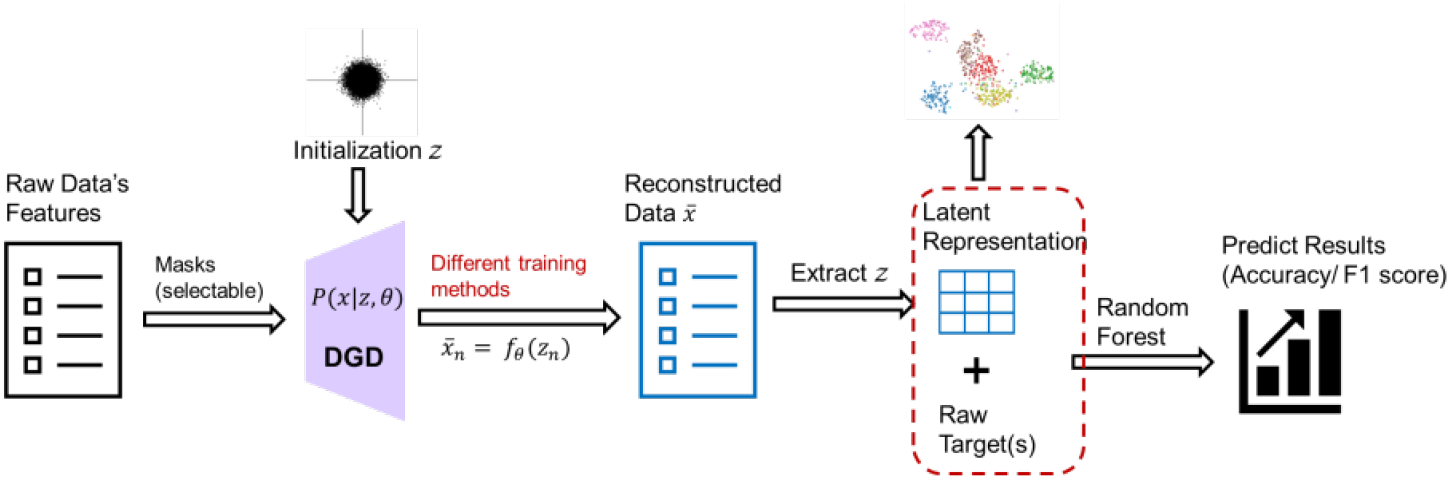
Overview of the DGD-based training and evaluation pipeline. Starts with raw data features, the DGD module initializes a latent variable z and generates reconstructed data through different training methods. From the reconstructed data, we extract latent representations and combine them with the raw targets. Finally, a Random Forest model is used to predict the results, evaluated by Accuracy or F1 score.

## D Optimization Hyperparameters for Different Methods and Datasets

Feature Batch Size = 5. Random Drop Rate = 0.3

**Table S1:**
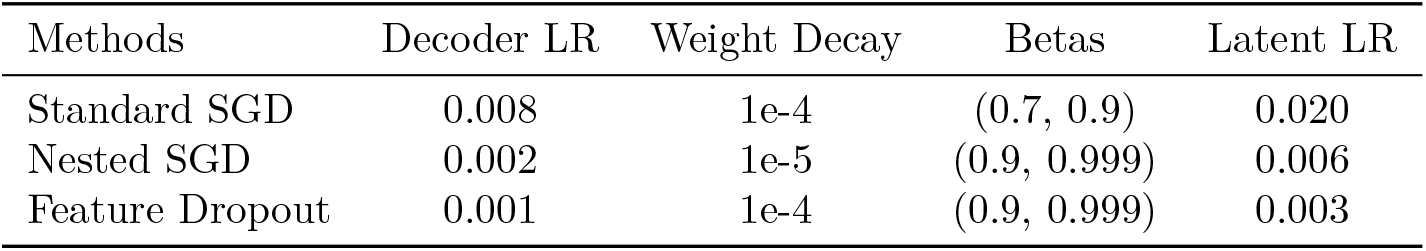
Adam optimizer parameters for Diabetes disease.

**Table S2:** Adam optimizer parameters for Thyroid Disease.

| Methods | Decoder LR | Weight Decay | Betas | Latent LR |
| --- | --- | --- | --- | --- |
| Standard SGD | 0.002 | 1e-5 | (0.9, 0.999) | 0.006 |
| Nested SGD | 0.002 | 1e-5 | (0.9, 0.999) | 0.006 |
| Feature Dropout | 0.002 | 1e-5 | (0.9, 0.999) | 0.005 |

**Table S3:**
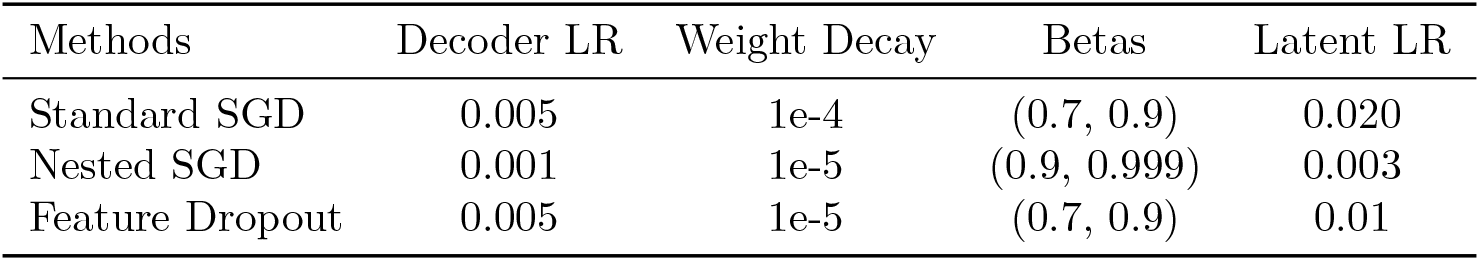
Adam optimizer parameters for Parkinson Telemonitoring.

| Methods | Decoder LR | Weight Decay | Betas | Latent LR |
| --- | --- | --- | --- | --- |
| Standard SGD | 0.005 | 1e-4 | (0.7, 0.9) | 0.020 |
| Nested SGD | 0.001 | 1e-5 | (0.9, 0.999) | 0.003 |
| Feature Dropout | 0.005 | 1e-5 | (0.7, 0.9) | 0.01 |

**Table S4:** Learning rate scheduler parameters (ReduceLROnPlateau) for each optimizer.

| Optimizer | Dataset | Mode | Factor | Patience |
| --- | --- | --- | --- | --- |
| Model optimizer | all | min | 0.5 | 4 |
| Latent z | all | min | 0.5 | 4 |
| Latent z val | all | min | 0.5 | 4 |
| Latent z test | all | min | 0.5 | 4 |

